# CT-FOCS: a novel method for inferring cell type-specific enhancer-promoter maps

**DOI:** 10.1101/707158

**Authors:** Tom Aharon Hait, Ran Elkon, Ron Shamir

**Affiliations:** The Blavatnik School of Computer Science, Tel Aviv University, Tel Aviv, 69978, Israel; Department of Human Molecular Genetics and Biochemistry, Sackler School of Medicine, Tel Aviv University, Tel Aviv, 69978, Israel; Sagol School of Neuroscience, Tel Aviv University, Tel Aviv, 69978, Israel

## Abstract

Spatiotemporal gene expression patterns are governed to a large extent by the activity of enhancer elements, which engage in physical contacts with their target genes. Identification of enhancer-promoter (EP) links that are functional only in a specific subset of cell types is a key challenge in understanding gene regulation. We introduce CT-FOCS, a statistical inference method that uses linear mixed effect models to infer EP links that show marked activity only in a single or a small subset of cell types out of a large panel of probed cell types. Analyzing 808 samples from FANTOM5, covering 472 cell lines, primary cells, and tissues, CT-FOCS inferred such EP links more accurately than recent state-of-the-art methods. Furthermore, we show that strictly cell type-specific EP links are very uncommon in the human genome.

## INTRODUCTION

Understanding the effect of the noncoding part of the genome on gene expression in specific cell types is a central genomic challenge (1). Cell identity is, to a large extent, determined by transcriptional programs driven by lineage-determining transcription factors (TFs; reviewed in (2)). TFs mostly bind to enhancer elements located distally from their target promoters (3). Furthermore, the expression of a gene can be regulated by different enhancers in different cell types. For example, TAL1 transcription is regulated by three enhancers, two of which are active in different cell types (HUVEC and K562)(4). To find enhancer-promoter (EP) links that are active in only very few cell types (hereafter, referred to as ct-links), one has to compare links across multiple and diverse cell types. 3D chromatin conformation data, which can identify ct-links, e.g., ChIA-PET (5), HiChIP (6) and Hi-C (7, 8), are still not available for many cell types and tissues (7–12). Consequently, there is a high need for computational methods that predict ct-links based on other data. A key resource for such prediction is large-scale epigenomic data, which are available for a variety of human cell types and tissues, and enable quantification of both enhancer and promoter activities.

A key challenge is to identify which of the numerous candidate EP links are actually (1) functional (or active) and (2) show their activity only in a specific small subset of cell types of interest. Ernst et al. (13) predicted ct-links based on correlated cell type-specific enhancer and promoter activity patterns from nine chromatin marks across nine cell types. Similarly, the Ripple method (14) predicted ct-links in five cell types. The cell-type specificity of the inferred EP links was quantified by comparison of their occurrence in other cell types. Additional methods that predicted EP links that are specifically active in a low number of cell types are IM-PET (15) and TargetFinder (16). All these methods used data of multiple chromatin marks and gene expression data for the studied cell types. The JEME algorithm finds global and cell type-active EP links (but not necessarily cell type-specific), using one to five different types of omics data (17). Each EP link reported by JEME is given a score for its tendency to be active in a given cell type. JEME reported an average of 4,183 active EP links per cell type, and many of these may show a broad activity profile. Fulco and Nasser et al. (18, 19) recently introduced the Activity-By-Contact (ABC) score for inferring cell type-specific functional EP links in 131 human biosamples with an average of 48,441 EP links per biosample. The ABC score was calculated using read counts of DNase Hypersensitive Site (DHS) and H3K27ac chromatin immunoprecipitation sequencing (ChIP–seq) at enhancer elements, and Hi-C contact frequency between enhancers and promoters.

Evidence of several sources suggests that while each cell type manifests tens of thousands of EP links, most of them are not unique and are shared across cell types. In a recently published compendium of EP chromatin interactions across 27 human cell types (20), the number of EP loops that were unique to a specific cell type was rather low (a median of 630 unique EP links, compared to a median of 31,250 total EP links per cell type) (**Methods**). In line with these numbers, comparing 3D genome architecture between neuronal progenitor cells (NPC) and mature neurons, Rajarajan et al. (21) identified 1,702 and 442 NPC- and neuron-specific chromatin loops linked to 386 and 385 genes, respectively. Similarly, the portion of EP links identified by JEME and ABC models in only one cell type was low (~30% and ~32%, respectively).

Here, we develop a novel statistical method for inferring ct-links from large-scale compendia of cell types measured by a single omics technique. We take advantage of linear mixed effect models to estimate cell-type activity coefficients based on replicates available for each cell type. We compared the results to those of extant methods in terms of concordance with experimentally derived chromatin interactions and cell specificity of gene expression.

## MATERIALS AND METHODS

### FANTOM5 and ENCODE data preprocessing

Please refer to the Supplemental Methods in the Supplemental Material for ‘FANTOM5 CAGE data preprocessing’ and ‘ENCODE DHS data preprocessing’ sections.

### CT-FOCS model Implementation

Our model for promoter *p* (**Fig. 1**) includes its *k* closest enhancers. The activity of the promoter across the *n* samples is denoted by the *n*-long vector *y*_*p*,_ and the activity level of the enhancers across the samples is summarized in the matrix *X_e_* of dimensions *n* × (*k* + 1), with the first column of ones for the intercept and the next k columns corresponding to the candidate enhancers. There are *C* < *n* cell types and each sample is labeled with a cell type. *k* = 10 was used.

**Figure 1.**
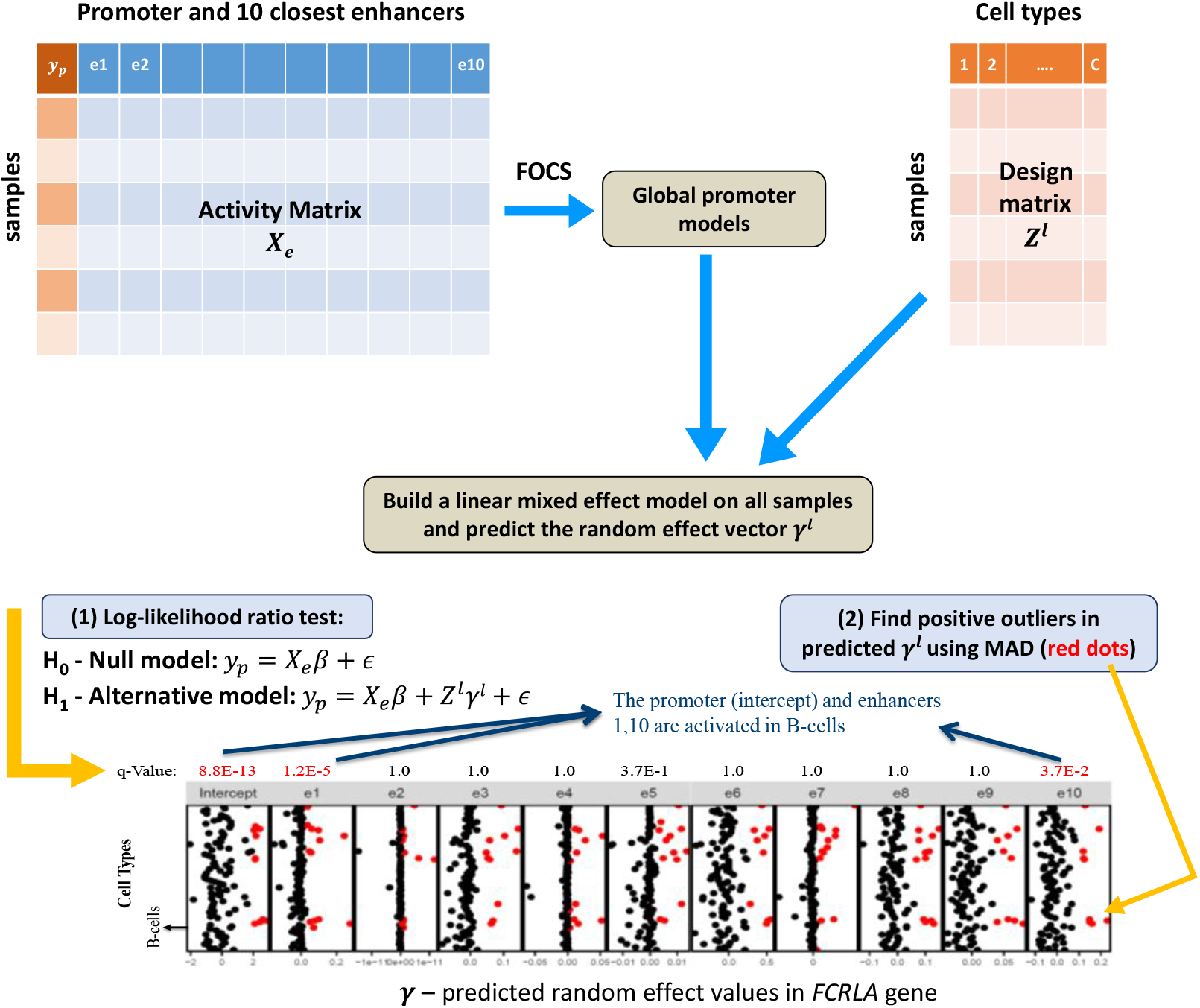
Outline of the CT-FOCS algorithm. Let *y_p_* denote the observed activity of promoter *p*, and *X_e_* be the activity matrix of the *k* = 10 closest enhancers to *p*. If *l* ∈ {1,…,*k* + 1} is one of the variables (enhancer or promoter, i.e. the intercept), then *Z^l^*[*i*, *j*] equals to *X_e_*[*i*, *j*] if sample *i* belongs to cell type *j* and 0 otherwise (see **Methods**). First, a robust global promoter model is inferred by applying the leave-cell-type-out cross validation step in FOCS (see Hait et al. 2018 for details). Second, a linear mixed effects model (LMM) is built on all samples using *y_p_*, *X_e_*, and *Z^l^*. The LMM includes the component *Z^l^γ^l^* where *γ^l^* is a vector of the predicted random effect values for each variable (i.e., enhancer or promoter) per cell type. Then, the algorithm performs two tests for every *l*: (1) log-likelihood ratio test (LRT) to compare the simple linear regression and the LMM model. The test is carried out eleven times (testing the 10 enhancers and the intercept). The p-values for these LRTs are adjusted for multiple testing (*q*-values). (2) The *γ^l^* values produced by the LMM are standardized using the Median Absolute Deviation (MAD) technique and positive outliers (red dots) are identified. A cell type-specific EP link (ct-link) is called if: (1) both enhancer and promoter (i.e., the intercept) have *q*-value <0.1 (marked in red), and (2) the enhancer and the promoter are found as positive outliers in the same cell type. In the *FCRLA* gene given as an example, the promoter *p* and enhancers *e*_1_, *e*_10_ are significant and are commonly found as positive outliers in B-cells. Therefore, E1p and E10p are called by CT-FOCS as B-cell-specific EP links.

To find ct-links based on the global links identified by FOCS, CT-FOCS starts with the full (that is, non-regularized) promoter model. We use the non-regularized promoter model as regularization reduces the overall model variance needed for making inferences. In principle, one could apply ordinary least squares regression with the cell types as additional coefficients to estimate cell type specificity. However, such models will perform poorly when the sample size is not much larger than the number of coefficients (e.g., in FANTOM5 we have 808 samples and a total of 483 coefficients: 472 cell types + k=10 enhancers + intercept). By using LMM, we can treat the cell type group level as a random effect coefficient, splitting the samples (replicates) based on their cell type of origin, at the cost of assuming a random effect distribution.

The application of an appropriate mixed effects model to the data depends on the distribution of the promoter and enhancer activities. We observed that FANTOM5 data have normal-like distribution and ENCODE data have zero-inflated negative binomial (ZINB) distribution (**Supplementary Fig. S1**). For FANTOM5, we applied regular linear mixed effect regression. For ENCODE, we applied generalized linear mixed effect regression (GLMM).

For each promoter, we defined a null model and *k* + 1 alternative models, each corresponding to a single random effect (i.e., random slope for enhancer or random intercept for the promoter). We defined the null model as the simple linear regression *y_p_* = *X_e_β* + *ϵ*, and each of the alternative models as the LMM model *y_p_* = *X_e_β* + *Z^l^γ^l^* + *ϵ*, where *X_e_β* is the fixed effect, *Z^l^γ^l^* is the random effect, and *ϵ* is a random error. *l* ∈ {1,…,*k* + 1} is one of the variables (enhancer or the intercept). *γ^l^* is a C-long vector of random effects to be predicted. *Z^l^* is a *nxC* design matrix that groups the samples by their cell types, namely:

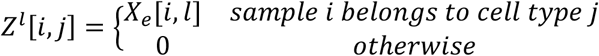

We applied a likelihood ratio test between the residuals of the *k* + 1 alternative models and the null model, and got *k* + 1 *p*-values. Such *p*-values were calculated for each of the |*P*| promoters, and corrected together for multiple testing using FDR (22), with the number of tests performed |*P*| · (*k* + 1).

Each predicted random effect vector 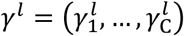 of the alternative models was normalized using the median absolute deviation (MAD), i.e., 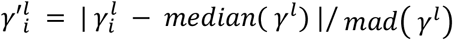, where *mad*(*γ^l^*) = *median*(|*γ^l^* – *median*(*γ^l^*)|) is calculated over all cell types together. If 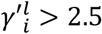 then enhancer *l* (or the promoter, if *l* = 1) was regarded as having an outlier activity in cell type *i*. We chose a moderately conservative MAD threshold, 2.5, as suggested in (23). We chose to use the MAD statistic since the mean and the standard deviation are known to be sensitive to outliers (23).

Finally, we defined cell type-specific EP links (abbreviated *ct-links*) as those that had: (1) significant random effect intercept of the promoter (P), (2) significant random effect slope of the enhancer (E), both with *q*-value < 0.1, and (3) E and P were identified as outliers in the same cell type according to the MAD criterion.

### MAD-FOCS model

MAD-FOCS takes the global EP links predicted by FOCS, before shrinkage (24). Then, for every global EP link, MAD-FOCS calculates the E and P median activity values across the multiple replicates per cell type. Last, it normalizes the median activities across cell types using the median absolute deviation (MAD) method. EP links are identified as ct-links in a certain cell type if both E and P are positive outliers in that cell type using MAD cutoff > 2.5.

### Filtered EP links sets

#### 1. CT-JEME model

JEME reports a classification score (between 0.3 to 1) for every EP link representing how active the EP link is in each cell type. To make a fair comparison between the predictions of CT-FOCS and JEME on the FANTOM5 dataset, we created a filtered set of JEME EP links called cell-type JEME (CT-JEME). For cell type j in FANTOM5 with *n* CT-FOCS ct-links, we chose the top *n* scoring EP links of JEME as the predicted CT-JEME solution for that cell type. For cell types in which JEME included a lower number of EP-links than CT-FOCS, we included all JEME’s EP links for that cell type in CT-JEME.

#### 2. CT-MAD-FOCS model

To allow a fair comparison between the predictions of CT-FOCS and MAD-FOCS, we created a filtered set of MAD-FOCS EP links called cell type MAD-FOCS (CT-MAD-FOCS), as decribed for CT-JEME above. We sorted the EP links by their *logEP* signal.

#### 3. CT-TargetFinder and CT-ABC models

Data for ABC model was taken from ftp://ftp.broadinstitute.org/outgoing/lincRNA/ABC/AllPredictions.AvgHiC.ABC0.015.minus150.ForABCPaperV3.txt.gz. Among the 131 biosamples analyzed in ABC, 75 were taken from ENCODE and Roadmap epigenomics consortia (25, 26) and 8 of them were also present in the CT-FOCS database and used for comparison (GM12878, HeLa-S3, K562, HCT-116, HepG2, A549 and H1-hESC). As for TargetFinder, we applied the program (https://github.com/shwhalen/targetfinder) on five cell types from ENCODE (GM12878, HeLa-S3, HUVEC, NHEK and K562) for which preprocessed multi omics data was available on the TargetFinder website, using as input candidate DHS sites representing enhancers and promoters from ENCODE DHS data. For each cell type in ENCODE with *n* CT-FOCS ct-links, we chose the top *n* scoring EP links of TargetFinder (by classification score) and of the ABC model (by ABC score) as the predicted solutions for that cell type for the two models. Statistics on the analyzed data are summarized in **Supplemental Table 1A**.

### External validation of predicted EP links using ChIA-PET, HiChIP and PCHi-C loops

We used 3D chromosome conformation capture loops to evaluate the performance of CT-FOCS and of other methods for EP linking. We downloaded ChIA-PET data of GM12878 cell line (GEO accession: GSE72816; ~100 bp resolution) assayed with *POLR2A* (12), HiChIP data of Jurkat, HCT-116, and K562 cell lines (GEO accession: GSE99519; 5 kb resolution) assayed with *YY1* (27), and PCHi-C data across 27 tissues (GEO accession: GSE86189; 5 kb resolution) (20).

Each loop identifies an interaction between two genomic intervals called its *anchors*. In ChlA-PET data, to focus on high confidence interactions, we filtered out loops with anchors’ width >5kb or overlapping anchors. Loop anchors were resized to 1kb (5kb in HiChIP and PCHi-C) intervals around the anchor’s center position. We filtered out loops crossing topologically associated domain (TAD) boundaries, as functional links are usually confined to TADs (9, 28–30). For this task, we downloaded 3,019 GM12878 TADs (31), which are largely conserved across cell types (8), and used them for filtering ChIA-PET and PCHi-C loops from all cell types.

To overcome the sparseness of the ChIA-PET loops, and the 8kb minimum distance between loop anchors (11, 12), we combined loops into two-step loop sets (**TLSs**) as follows: for every reference loop, x, its TLS is defined as the set of anchors of all loops that overlap with at least one of x’s anchors by at least 250 bp (**Fig. 2A**). We used the igraph R package (32) for this analysis.

**Figure 2.**
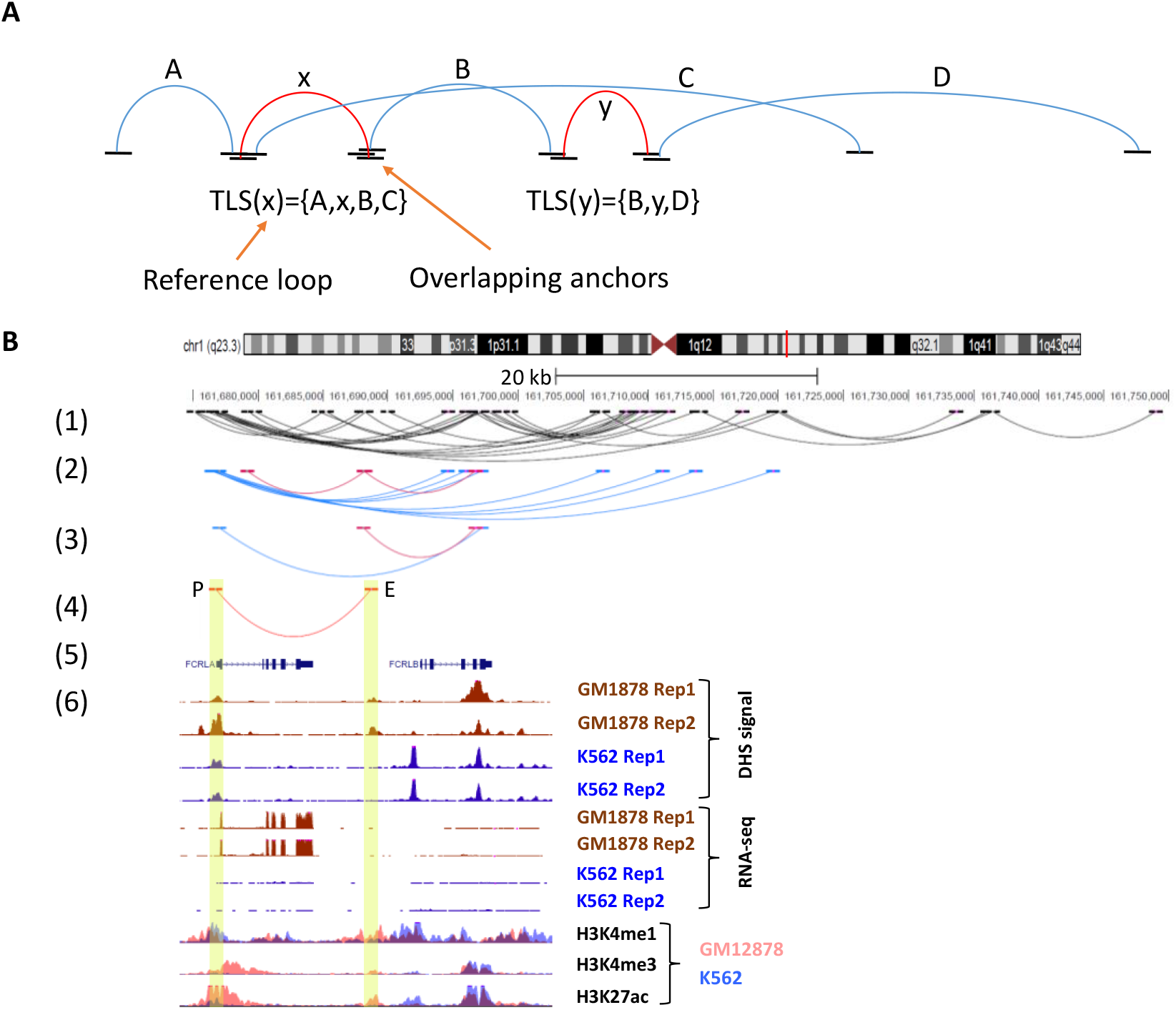
ChIA-PET TLSs support predicted ct-links. The two-step connected loop set (TLS) of reference loop x is defined as the set of all loops that have an anchor overlapping one of x’s anchors including loop x. (A) Examples of TLSs. Loop x’s anchors overlap with at least one of the anchors of loops A, B, and C, and, therefore, the TLS of x is composed of loops A, x, B, C. Similarly, the TLS of y is composed of loops B, y, and D. (B) (1) A 70kb region of Chromosome 1 showing ChIA-PET loops detected in cell type GM12878. (4) A ct-link predicted by CT-FOCS. (2) The same region showing only loops that have anchors overlapping the anchors of the ct-link. Pink: loops overlapping the enhancer; blue: loops overlapping the promoter. (3) A TLS that supports the predicted ct-link. The ct-link in (4) is validated by the TLS, but not by any single ChIA-PET loop. (5) Gene annotations. (6) Gene expression (RNA-seq) and epigenetics signals (DHS-seq and selected histone modifications) for the region. Tracks are shown using UCSC Genome Browser for data from GM12878 and K562 cell lines. The data supports the link in GM12878 but not in K562.

To evaluate if a ct-link is confirmed by the ChIA-PET data, we checked if both the enhancer and the promoter fall in the same TLS. Specifically, we defined 1kb genomic intervals (±500 bp upstream/downstream; 5kb genomic intervals: ±2.5kb upstream/downstream in HiChIP and PCHi-C) for the promoters (relative to the center position; relative to the TSS in FANTOM5 dataset) and the enhancers (relative to the enhancer’s center position) as their genomic positions. Both inter- and intra-TAD predicted EP-links were included in the validation. An EP link was considered supported by a TLS if the genomic intervals of both its promoter and enhancer overlapped different anchors from the same TLS **(Fig. 2B** and **Supplemental Fig S2)**. We used randomization in order to test the significance of the total number of supported EP links by ChIA-PET single loops. We denoted that number by *N_t_*. We performed the test as follows: (1) For each predicted EP link, we randomly matched a control EP link, taken from the set of all possible EP pairs that lie within 9,274 GM12878 TADs from Rao et al. (8), with similar linear distance between E and P center positions. We restricted the matching to the same chromosome in order to account for chromosome-specific epigenetic state (33). The matching was done using MatchIt R package (method=‘nearest’, distance=‘logit’, replace=‘FALSE’) (34). This way, the final set of matched control EP links had the same set of linear interaction distances as the original EP links. (2) We counted *N_r_*, the number of control EP links that were supported by ChIA-PET single loops. We repeated this procedure for 1,000 times. The empirical *p*-value was 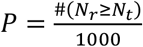, or *P*<0.001 if the numerator was zero. A similar empirical *p*-value was computed for the validation rate obtained by using single loops and TLSs.

We used the following formula to calculate the GM12878 ChIA-PET TLS support ratio:

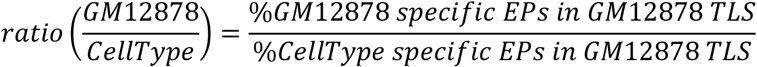

### Calling cell-type specific active EP loops reported in a capture Hi-C study

We wished to identify cell-type specific EP links reported in capture Hi-C data (20). We downloaded 906,721 promoter-other (PO) capture Hi-C loops generated across 27 tissues (GEO accession: GSE86189) (20). These loops involve a known gene’s promoter and a non-promoter region, which may be an enhancer. To define a set of strictly ct-specific loops, we retained PO loops that were detected in exactly one cell type. We set the PO anchors to 1kb intervals around their center positions. This analysis detected a median of 630 EP loops that were unique to a specific cell type.

To call promoter and enhancer regions, we downloaded 474,004 enhancer and 33,086 promoter regions predicted by a 15-state ChromHMM model on Roadmap epigenetic data across 127 tissues (https://personal.broadinstitute.org/meuleman/reg2map/HoneyBadger2-intersect_release/DNase/p10/enh/15/state_calls.RData; https://personal.broadinstitute.org/meuleman/reg2map/HoneyBadger2-intersect_release/DNase/p10/prom/15/state_calls.RData) (26). We kept the enhancers of state Enh or EnhG (genic enhancers) in any of 127 Roadmap tissues. Similarly, we kept the promoters of state TssA (active TSS) or TssAFlnk (Flanking Active TSS). Then, we resized each region to a 1kb interval around its center position. We called the resulting sets active promoters and enhancers. A retained PO loop whose P and O anchors had at least 250 bp overlap with active ChromHMM promoter and enhancer, respectively, was considered as cell type-specific active EP loop.

### Cell type specificity score

We quantified the intensity of an EP link in a given sample by log_2_ *a* + log_2_ *b* where *a* and *b* are the enhancer and promoter activities in that sample. The *EP signal* of the link for a particular cell type is the average of the signal across the samples from that cell type. Define *x_c_* = (*x*_*c*1_,…,*x_cn_*) as the vector of signals in cell type *c*, where *n* is the total number of EP links discovered in cell type *c*, and define *d_c,i_* as the Euclidean distance between the vectors of cell types *c* and *i*, both with the same EP links from cell type *c*. Following the definition of (35), the *specificity score* of EP links predicted in cell type *c* is:

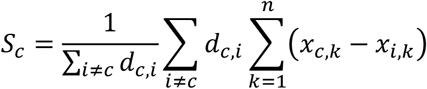

Similarly, cell-type specificity can be computed for the expression values of the genes annotated with EP links, or on the overrepresentation factors of TFs found at enhancers and promoters.

### Statistical methods, visualization and tools

All computational analyses and visualizations were done using the R statistical language environment (36). To correct for multiple testing we used the p.adjust() function (method=‘BY’). We used ‘GenomicRanges’ package (37) for finding overlaps between genomic intervals. We used ‘rtracklayer’ (38) and ‘GenomicInteractions’ (39) packages to import/export genomic positions. Linear mixed effect regression models were created using lme R function from nlme package (40). Generalized linear mixed effect with zero inflated negative binomial models were created using glmmTMB R function from glmmTMB package (41). Counting reads in genomic intervals was done using BEDTools (42). Graphs were created using graphics (36), ggplot2 (43), gplots (44), ComplexHeatmap (45), and the UCSC Genome Browser (https://genome.ucsc.edu/).

## RESULTS

### The CT-FOCS algorithm

We developed a novel method called CT-FOCS (*C*ell *T*ype FOCS) for inferring ct-links. The method utilizes a single type of omics data (e.g., CAGE or DHS) from large-scale datasets.

The input to CT-FOCS is enhancer and promoter activity profiles for a set of cell types. The output is the set of ct-links called for each cell type. Note that the enhancers or promoters involved in ct-links can be broadly active separately. In contrast to methods that seek global correlations between the activity profiles of enhancers and promoters, the aspect emphasized and detected by CT-FOCS is the specificity of the link between the two elements: that is, links reported by CT-FOCS highlight the few cell types in which the enhancer and promoter are predicted to functionally interact.

CT-FOCS builds on FOCS (24), which discovers global EP links showing correlated enhancer and promoter activity patterns across many samples. FOCS performs linear regression on the levels of the 10 enhancers that are closest to the target promoter, followed by two non-parametric statistical tests for producing initial promoter models, and regularization to retrieve the most informative enhancers per promoter model. CT-FOCS starts with the full (non-regularized) FOCS promoter model (**Methods)**, and uses a linear mixed effect model (LMM), utilizing groups of replicates available for each cell type to adjust a distinct regression curve per cell-type group in one promoter model (**Fig. 1; Methods**). We call a ct-link in a certain cell type if it meets the following criteria: (1) both the enhancer (E) and the promoter (P) show markedly positive activity levels in that cell type compared to other cell types, and (2) both P and E have significantly high random effect coefficients, reflecting an advantage of the LMM over the global FOCS model (**Methods**). The second criterion increases our confidence that the high activity detected by the first is specific to this cell type.

To demonstrate the difference between the linear and LMM predictions, **Supplemental Fig S3** shows, for the same promoter (P), two links involving distinct enhancers (E1 and E2), one predicted by CT-FOCS (E1P) and the other (E2P) by FOCS. The link between E1 and P is active only in neurons, while the link between E2 and P is active over a wider range of cell types of distinct lineages (amniotic membrane cells, whole blood cells, fibroblasts, endothelial cells and preadipocytes).

We applied CT-FOCS on FANTOM5 cap analysis of gene expression (CAGE) profiles, which include 808 samples from 225 cell lines, 157 primary cells, and 90 tissues (46) (**Methods**). CAGE quantifies the activity of both enhancers and promoters, and overall this dataset covers 42,656 enhancers and 24,048 promoters (mapped to 20,597 Ensembl protein-coding genes). For some analyses, we also applied CT-FOCS to ENCODE’s DNase Hypersensitive Site (DHS) profiles (25, 47), which cover 106 cell types, each with typically 2 replicates. This dataset includes measurements for 36,056 promoters (mapped to 13,464 Ensembl protein-coding genes) and 658,231 putative enhancers (**Methods**). Unlike the FANTOM5 dataset, open genomic regions identified by DHS do not necessarily mark functionally active enhancers and promoters. Thus, EP maps inferred using the ENCODE dataset may be less reliable, and we focus our analyses mainly on the FANTOM5 dataset.

Overall, CT-FOCS identified 195,232 ct-links in FANTOM5 dataset **(Table 1)**, with an average of 414 ct-links per cell type (median 594, **Table 1; Supplemental Fig S4A**). These results are in line with the low number of ct-links observed experimentally by the abovementioned studies, including for NPC and neurons (21, 48), and further indicate that the EP links specific to a cell type constitute only a small portion of the EP links that are active in it. The EP links called by CT-FOCS were on average shared across 2.5 cell types (**Supplemental Fig S4B**). CT-FOCS predicted both proximal and distal interactions, with an average EP distance of ~160kb (median ~110kb; **Supplemental Fig S4C**). The complete set of predicted ct-links for each cell type is available at http://acgt.cs.tau.ac.il/ct-focs.

**Table 1.** Statistics on the number of CT-FOCS predictions per cell type

| Dataset | Avg. ct- links | Avg. enhancers | Avg. promoters | Avg. genes* | Tot. ct-links | Cell type with maximum ct-links |
| --- | --- | --- | --- | --- | --- | --- |
| FANTOM5 | 414 | 318 | 146 | 134 | 195,232 | Temporal lobe (13,354) |
| ENCODE | 167 | 158 | 86 | 82 | 17,672 | Caco-2 (1,572) |

(\*) Ensembl protein-coding genes

Since EP links are expected to function mostly within topologically associated domains (TADs) (49, 50), we next tested if ct-links detected by CT-FOCS are enriched for intra-TAD genomic intervals. As TADS are largely cell-type invariant (8), we used for these tests the 9,274 TADs reported by Rao et al. in GM12878 (8). Indeed, comparison with randomly matched EP links demonstrated that predicted ct-links tend to lie within TADs (**Supplemental Fig S5**).

### Inferred ct-links correlate with cell type-specific gene expression

To evaluate the specificity of the CT-FOCS predictions, we compared the activity of the set of ct-links inferred for a particular cell type with their activity in all other cell types. We defined the activity of an EP link in a cell type as the logarithm of the product of the enhancer and promoter activities in that cell type. We used these measures to compute the cell-type specificity for the set of ct-links detected in each cell type, using a score akin to (35) (**Methods**). As an example, CT-FOCS called 340 ct-links on the GM12878 lymphoblastoid cell line. We scored the cell-type specificity of these 340 ct-links for each cell type. Reassuringly, GM12878 was the top scoring cell type, and other high scoring cell types were enriched for related lymphocyte cells (other B-cells and T-cells; **Fig. 3A, C**). GM12878 was also ranked first in cell type-specificity scores calculated separately for the promoters and enhancers of these 340 ct-links (**Supplemental Fig S6;** see **Supplemental Fig S7** for additional example for neurons cells).

**Figure 3.**
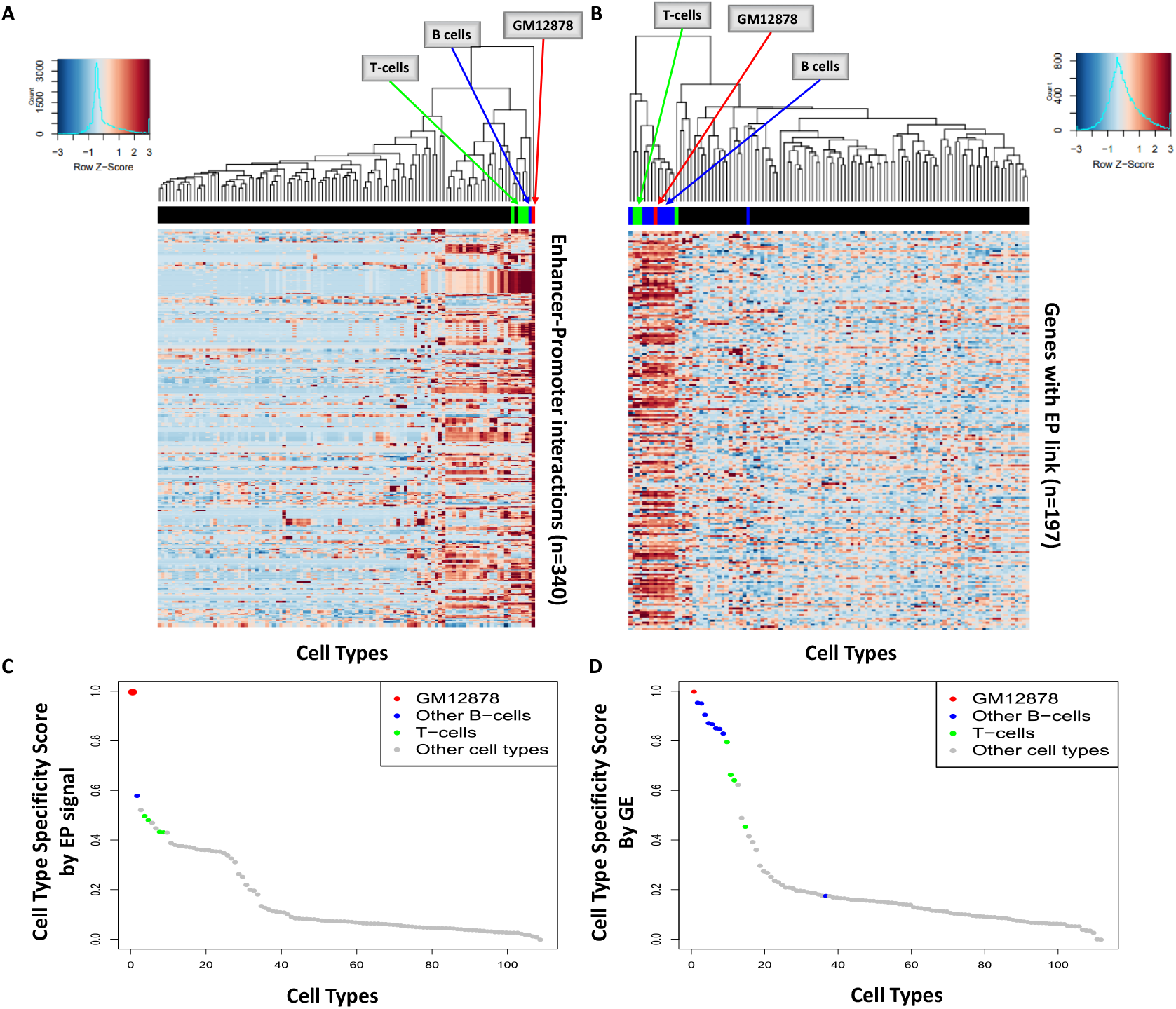
Specificity of ct-links predicted for GM12878 cell line. **(A)** Heatmap of EP signals for 340 ct-links predicted on GM12878 cells. Rows – EP links, columns – cell types, color – z-score of EP signal. Cell types related to lymphocytes (B/T-cells) are highlighted in color. **(B)** Heatmap of gene expression (GE) for 197 genes involved in the predicted ct-links. Rows – genes, columns – cell types, color – z-score of GE. **(C)** Cell type specificity scores based on the EP signals. **(D)** Cell type specificity scores based on expression for the gene set in B (Methods). In A and C, 109 cell types with at least 3 replicates are included in the analysis; in B and D, 112 cell types with ENCODE GE data are included (51).

Next, we examined how the effect of ct-links is reflected by cell-type specific expression of the linked genes (**Methods**). The 340 ct-links called by CT-FOCS in GM12878 involve 197 genes. We examined their expression profiles over 112 cell types using an independent gene expression (GE) dataset (51). In this analysis, we now scored each of the 112 cell types for the specificity in the expression of these 197 genes. Notably, here too, the lymphocyte group (B- and T-cells) showed the highest expression levels (**Fig. 3B**) with GM12878 ranking first by GE specificity (**Fig. 3D**). Overall, these results show that for GM12878, the ct-links predicted by CT-FOCS based on CAGE data are correlated with lymphocyte-specific GE programs.

### Comparison of CT-FOCS to other methods

We compared CT-FOCS predictions on the FANTOM5 dataset with those made by four alternative methods: (1) JEME (17), which predicts EP links that are active in a particular cell type but are not necessarily cell type-specific. (2) A naive variant of FOCS, which takes the shrunken promoter models from FOCS, and predicts ct-links by detecting cell types in which the promoter and any of the model’s enhancers show exceptionally high activity, based on the median absolute deviation (MAD) index. We call this variant MAD-FOCS (**Methods**). (3–4) Variants of JEME and MAD-FOCS with filtering of the reported links to produce sets of cell-type specific links of the same size as the ones detected by CT-FOCS (**Methods**). We call these variants cell-type-JEME (CT-JEME) and cell-type-MAD-FOCS (CT-MAD-FOCS), respectively.

**Supplemental Fig S4** shows basic properties of the solutions provided by the five methods. EP links predicted by JEME and MAD-FOCS were, on average, shared across 11 and 12 cell types (median=3 and 13 respectively; **Supplemental Fig S4B**). In contrast, the CT-FOCS, CT-MAD-FOCS and CT-JEME EP links were, on average, shared across <4 cell types (Median=2, 2 and 1, respectively), demonstrating that they identified EP links that are more specific. The same number of predicted links allows fair comparison between CT-FOCS, CT-MAD-FOCS and CT-JEME.

Next, we calculated cell-type specificity scores for the EP links called by CT-FOCS, CT-MAD-FOCS and CT-JEME on the 274 FANTOM5 cell types. For each cell type, we used the ct-links called on it to compare the specificity score of each cell type and ranked the scores. We expect the given cell type to score the top. In this analysis, CT-MAD-FOCS and CT-FOCS performed similarly, and significantly better than CT-JEME (**Supplementary Fig. S8A**). In terms of GE of the genes associated with the EP links, examining the four cell types that were present in both FANTOM5 and the independent GE data of 112 cell types of (51)(GM12878, K562, HepG2 and MCF-7), CT-FOCS was the only method which ranked 1^st^ all the four cell type (**Supplementary Fig. S8B**). Overall, these three methods seem to capture ct-links with highly specific EP and GE signals.

Next we ranked the cell types according to cell-type specificity scores obtained when considering separately the signals of the linked enhancers and promoters. The median rank of the enhancers was 1^st^ by all methods, possibly because enhancers tend to be cell type specific. However, the median rank of CT-JEME’s promoters was 23^rd^ while it was 1^st^ for CT-FOCS and CT-MAD-FOCS. The low ranks of CT-JEME’s linked promoters can explain why its predicted ct-links ranked lower compared to CT-FOCS and CT-MAD-FOCS.

Last, we compared the CT-FOCS predictions on ENCODE’s DHS dataset with those obtained by six other methods: (1–2) CT-MAD-FOCS and MAD-FOCS. (3) TargetFinder (16), which predicts EP links based on features in enhancer, promoter and the window between them using GradientBoosting trees. (4) ABC score model (18, 19), which inferred cell type-specific functional EP links in 131 human biosamples. (5–6) Variants of TargetFinder and ABC model having a similar number of predictions as CT-FOCS (**Methods**). We call these variants CT-TargetFinder and CT-ABC, respectively. Note that the comparisons on the FANTOM5 data were done on 274 cell types that had at least 50 predicted EP links in all methods. On the ENCODE dataset, the comparisons were done only on 5-10 cell types (**Methods**). Overall, CT-FOCS, CT-MAD-FOCS and ABC ranked first most cell types by specificity of the ct-link signals, better than the other methods. On the basis of GE specificity, CT-FOCS, ABC and CT-ABC ranked first most cell types, better than the other methods (**Supplemental Table 1B**).

### Introducing two-step connected loop sets in 3D chromatin conformation assays for evaluation of ct-links

We validated the ct-links predicted on GM12878 using both *POLR2A*-mediated ChIA-PET loops and promoter-capture (PC) Hi-C loops recorded in this cell line (12, 20). The direct way to validate a predicted ct-link is to check whether the E and P regions overlap the two anchors of the same loop. However, as loops indicate 3D proximity of their anchors, overlapping anchors of different loops indicate proximity of their other anchors as well (52, 53). Furthermore, ct-links that span a linear distance of < 20kb, where ChIA-PET loops may perform poorly (54), may not be supported by that assay. Thus, for the validation, we broadened the set of anchors that are considered to be proximal as follows: We define the two-step connected loop set (**TLS**) of a loop as the set of anchors of all loops that overlap with at least one of its anchors (**Fig. 2A**). We consider a predicted ct-link as validated if its enhancer and promoter regions overlap different anchors from the same TLS (**Fig. 2B;** see **Supplemental Fig S2** for an additional example**; Methods**).

Out of the 340 ct-links inferred by CT-FOCS in GM12878, 10% were supported by ChIA-PET single loops, and 33% were supported by TLSs. For PCHi-C loops, the numbers were 7.6% and 15%, respectively. Although these rates might seem low, in the next section we show that most methods predicting EP links have a low support from 3D conformation data. To test the significance of the observed validation rate, we generated random sets of 340 intra-TAD links having the same linear distances between E and P regions as the ct-links predicted by CT-FOCS (**Methods**). In 1,000 random sets, TLSs supported, on average, 9.4% (32 out of 340) and at most 14% (46 out of 340) (**Supplemental Fig S9A**), and the number of predicted ct-links supported by ChIA-PET data was significant with P<0.001. Similar significance was achieved when validating the predicted ct-links directly against single loops (**Supplemental Fig S9C)**. The same tests for PCHi-C loops gave an average overlap of matched random loops with PCHi-C TLSs of 8.5% (29 out of 340) and at most 12.4% (42 out of 340), with P=0.003 for TLS (**Supplemental Fig 9B**, P=0.048 for single loops; **Supplemental Fig S9D**).

### Validating predicted links by 3D conformation data

We compared the links predicted by CT-FOCS, CT-JEME and CT-MAD-FOCS to experimentally measured 3D chromatin loops, defined as the positive set. We chose the CT versions of these algorithms, which make the same number of calls, in order to allow fair comparison. In GM12878, CT-JEME achieved the best precision (21%) followed by CT-MAD-FOCS (19%) and CT-FOCS (10%). In K562, CT-FOCS achieved the best precision (17.5%) followed by CT-MAD-FOCS (14%) and CT-JEME (3.45%). The low precision shows that single loops do not support the majority of the links predicted by any method.

Repeating the comparison using TLSs instead of single loops resulted in 2-3 fold increase in precision compared to single loop validation in all methods. On GM12878 loops, precision was 54%, 50% and 30% in CT-JEME, CT-MAD-FOCS and CT-FOCS, respectively. On K562 loops, precision was 33%, 28% and 22% in CT-FOCS, CT-MAD-FOCS and CT-JEME, respectively. Again, the precision obtained by TLS validation for all methods was still low.

We repeated the same analysis on the ENCODE DHS dataset, comparing CT-FOCS to CT-TargetFinder and CT-ABC. Here, CT-FOCS performed markedly better in validation based on single loops and on TLSs. For example, on GM12878 with single-loop validation, CT-FOCS achieved 31% precision while CT-TargetFinder and CT-ABC model achieved 10% and 13%, respectively. With TLS validation, CT-FOCS had 66% precision while CT-TargetFinder and CT-ABC model achieved 30% and 47%, respectively. Similarly, on K562 with single loop validation, CT-FOCS had 54% precision, CT-ABC 30% and CT-TargetFinder 1.4 %. With TLS validation, CT-FOCS had 74% precision, CT-ABC 43% and CT-TargetFinder 3.7%.

Overall, ct-links predicted by all methods had low support from 3D chromatin loops. CT-FOCS tended to achieve higher precision than the other tested methods.

### Assessing cell type-specificity via 3D experimental loops

As an additional test, we checked to what extent ct-links called on different cell types are supported by GM12878’s POLR2A ChIA-PET TLS loops. If ct-links called on GM12878 are indeed highly specific, we expect GM12878 to show the highest support rate in this analysis. To quantify this, we defined for each cell type, the logarithm of the ratio between the validation rate observed in GM12878 and the validation rate observed for that cell type. For most cell types we expect to obtain values>0. Indeed, CT-FOCS ct-links predicted for GM12878 showed significantly higher support rate compared to the ct-links that were predicted in most other cell types (median log2(ratio) ~1.7; **Fig. 4A**). Moreover, the six cell types that showed higher validation rate than GM12878 (that is, had log2(ratio)<0; **Fig. 4A: CT-FOCS boxplot**) were all biologically related to GM12878 (e.g., B cell line and Burkitt’s lymphoma cell line). CT-MAD-FOCS and MAD-FOCS performance was significantly lower (median log2(ratio) ~1.1), followed by CT-JEME (~0.7) and JEME (~0.6). Note that in this analysis too, the comparisons between CT-FOCS, CT-MAD-FOCS and CT-JEME are more proper, since these methods have a similar number of predictions per cell type (and thus, comparable recall). The results for MAD-FOCS and JEME are added only for reference. We obtained similar results when validating against ChIA-PET single loops (**Fig. 4B**), and when using HiChIP from K562 (**Fig. 4C**). When using PCHi-C, HiChlP and ChlA-PET for eight individual tissues, CT-FOCS performed best overall (**Fig. 4D** and **Supplemental Table 2**).

**Figure 4.**
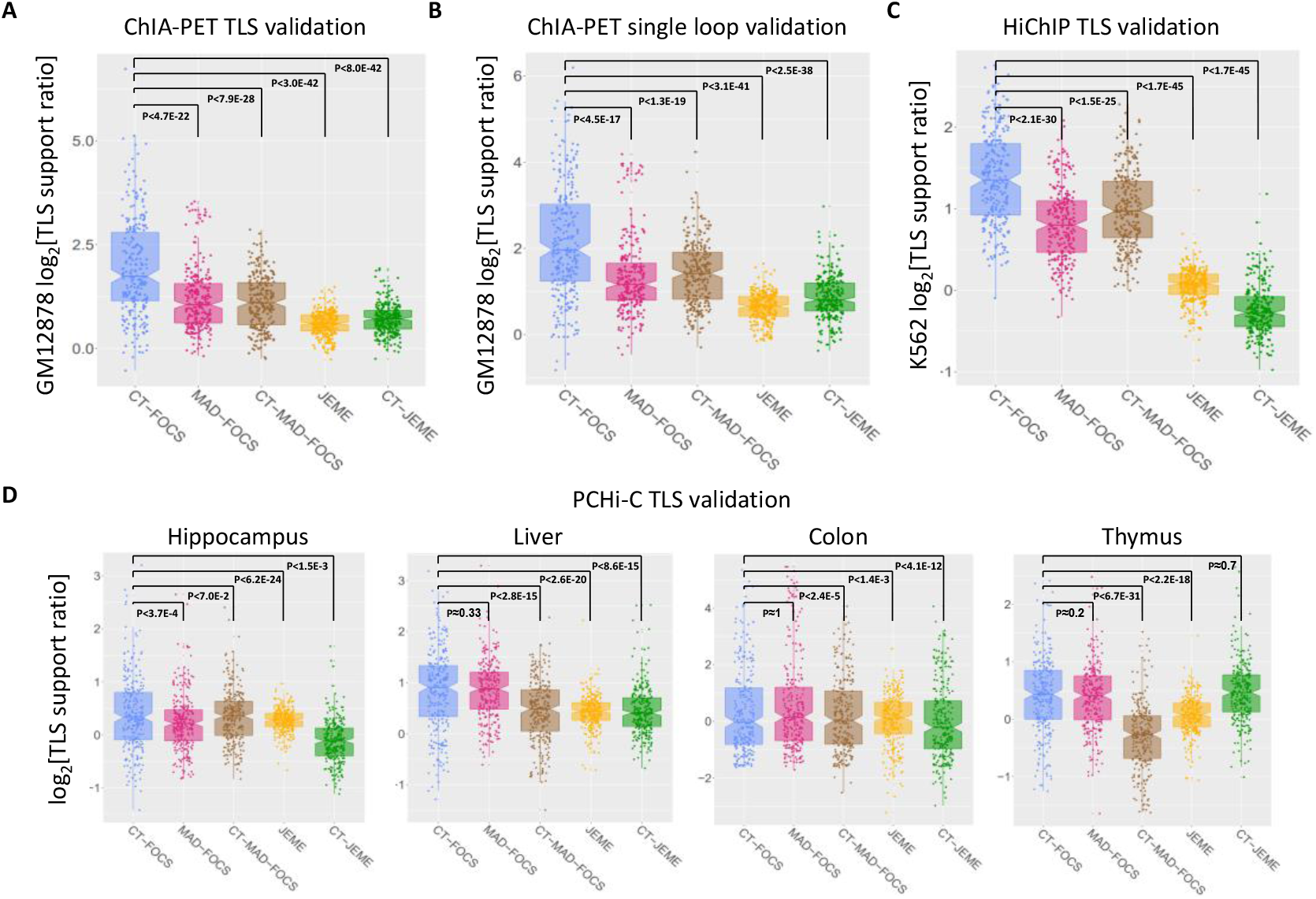
The particularity of each algorithm’s predictions as measured by ChIA-PET, HiChIP, and PCHi-C assays. **(A-B)** Each algorithm was applied to each cell type, and the predicted links were benchmarked against GM12878 ChIA-PET loops and TLSs. Comparison included 265 FANTOM5 cell types that had at least 50 predicted EP links in CT-FOCS, MAD-FOCS, CT-MAD-FOCS, JEME and CT-JEME. The plots show for each cell type the distribution of the ratios between the percentage of predicted EP links on GM12878 that had GM12878 ChIA-PET support and the percentage of predicted links in that cell type that had GM12878 ChIA-PET support (**Methods**). **(A)** ChIA-PET TLS support. **(B)** ChIA-PET single loop support. **(C)** The same analysis as in (A) for K562 cell line compared to TLSs derived from K562 HiChIP assay. **(D)** The same analysis as in (A) but here using TLSs derived from PCHi-C in four additional cell types and tissues. All comparisons are summarized in **Supplemental Table 2**. *p*-values are based on one sided Wilcoxon paired test.

We repeated the analysis on CT-FOCS, CT-MAD-FOCS, CT-TargetFinder and CT-ABC on ENCODE dataset (**Supplemental Table 1C)**. lnterestingly, CT-MAD-FOCS obtained the highest precision and TLS support on GM12878. On K562 cell type, across all cell types, all methods had low TLS support *(log_2_(ratío) ≈* 0). Note, however, that the number of cell types compared was very low (5-10 cell types, compared to 265 for FANTOM5), so these results are anecdotal. Overall, on FANTOM5 dataset, the particularity of the links of CT-FOCS was higher than those of CT-MAD-FOCS and CT-JEME.

## DISCUSSION

In this study we investigated the cell type-specificity of predicted EP links by state-of-the-art methods and introduced CT-FOCS, a novel method for inferring cell type-specific EP links (ct-links) based on activity patterns derived from a single type of omics data. We applied CT-FOCS on CAGE profiles from FANTOM5 (46). The resulting compendium of 195,232 ct-links for 472 cell types and the program are available for use at http://acgt.cs.tau.ac.il/ct-focs and enable further inquiry on gene regulation.

We compared the cell type-specificity of links predicted by each method on FANTOM5 data. We computed cell type-specificity scores by using either EP signals or target gene expression (**Supplementary Fig. S8**; **Supplemental Table 1A-B**; **Methods**). We found that CT-FOCS and CT-MAD-FOCS achieve similar and slightly better cell type-specificity ranks compared to CT-JEME on EP signal and target GE (**Supplementary Fig. S8**). Additionally, we introduced the two-step loop set (**TLS**) support ratio for benchmarking predicted ct-links against chromatin-interaction datasets (**Fig. 4**; **Supplemental Tables 1-2**; **Methods**). Using this criterion, we showed that the cell-type particularity of ct-links predicted by CT-FOCS was significantly higher than those of CT-JEME and CT-MAD-FOCS in 5-6 out of 8 examined cell types with available 3D conformation data (**Fig. 4; Supplemental Table 2**).

Several comments are in order regarding our inferred ct-links. First, a common naïve practice is to map enhancers to their nearest gene. Among the CT-FOCS predicted EP links, on average per cell type, only ~10% of the enhancers mapped to the nearest gene. While this proportion is lower than observed in previous reports (~74% in FOCS and ~40% in FANTOM5 (46)), it may have been affected by the relatively low number of FANTOM5 reported enhancers (~43k) due to lower sensitivity of detecting enhancers using CAGE data (55). FANTOM5 enhancers tend to be located in intergenic regions, possibly reducing the correlation of the enhancers with the nearest gene, which is more apparent for intragenic enhancers located within introns of the target genes. As a result, fewer EP links are identified using correlation-based techniques (e.g., linear regression). On the other hand, low-distance links were reported to have poor validation results in ChIA-PET and Hi-C 3D loops and eQTL data (17). Second, an average of ~60% of the predicted ct-links involve intronic enhancers, similar to the report by FOCS (70%). Third, the average number of predicted ct-links per cell type was rather modest: 414 in FANTOM5 (**Table 1**). This relatively low number is in line with the small number of ct-links reported previously in experiments on NPC, neuron, and K562 cells (21, 48), suggesting that only a small portion of the EP links that are active in a cell type are specific for it. Fourth, on average, per cell type, promoters were linked by ct-links to ~2 (and a maximum of 9) enhancers.

In terms of methodology, CT-FOCS uses linear mixed models to account for two effects. The first is the joint contribution of multiple enhancers to the promoter activity, which was shown to predict gene expression more accurately than to pairwise enhancer-gene correlations (17). The second is the contribution of distinct cell-type groups to promoter activity. By considering the cell-type effect, prediction of promoter activity can be done separately for each cell-type group. Thus, the estimated regression coefficient will not be the same for all samples but rather adjusted according to their cell type. In this way, ct-links are inferred based on the difference in the regression coefficients estimated for different cell type groups.

FOCS predictions are based on leave-cell-type-out cross validation. As such, by design, it cannot infer models that are strictly cell type-specific (24) (that is, EP pairs that are active in only one specific cell type and have completely null activity in all the rest). As CT-FOCS is built upon FOCS predictions, this limitation is true for CT-FOCS predictions as well. However, we confirmed in the broad epigenomic datasets that we analyzed, that cases in which an enhancer is active in only one cell type are very rare (**Supplemental Results** – ‘ Loops involving enhancers active in a single cell type’). Nevertheless, CT-FOCS EP links show very high cell-type specificity: they were shared, on average, by not more than three cell types (**Supplemental Fig S4B**), and >44% of them were called in a single cell type. The links identified by CT-FOCS correspond to much more prevalent (and therefore, biologically more relevant) cases, in which an enhancer shows activity in several (typically, highly related) cell types, but its impact on the activity on the target promoter is markedly more prominent in one or very few of them.

A limitation of CT-FOCS is that it considers only the ten closest enhancers to each promoter when building the models. A possible future improvement to CT-FOCS is to include all enhancers within a window of 1Mb around each promoter, e.g., by using Bayesian hierarchical models, considering possible confounders and a-priori information such as ChIA-PET and PCHi-C loops and eQTLs.

CT-FOCS can be useful for multiple genomic inquiries. For example, it can improve identification of known and novel cell type-specific TFs and enhance our understanding of key transcriptional cascades that determine cell fate decisions. Furthermore, the integration of protein-protein interactions (PPIs) with TF identification in predicted ct-links may help identify cell type-specific PPI modules (56). These modules may contain additional new proteins (e.g., co-factors and proteins that are part of the mediator complex) that shape the 3D chromatin in a cell type-specific manner. Overall, the new method we introduced and the compendium of ct-links can advance our understanding of cell type-specific genome regulation.

## DATA AVAILABILITY

CT-FOCS predicted ct-links and data are freely available at http://acgt.cs.tau.ac.il/ct-focs. Our database also contains ct-link predictions for 73 Roadmap Epigenomics cell types (26). CT-FOCS source code is freely available at http://acgt.cs.tau.ac.il/ct-focs, GitHub (https://github.com/Shamir-Lab/CT-FOCS).

