## Supplemental Material for "CT-FOCS: a novel method for inferring cell type-specific enhancer-promoter maps"

**Outline:**

**Supplemental Results**

**Supplemental Tables**

**Supplemental Figures**

**Supplemental Methods**

**Supplemental References**

**Supplemental Results**

**Loops involving enhancers active in a single cell type**

We wished to evaluate the prevalence of experimentally detected EP links that involve enhancers that are active only in one single cell type. We performed the following analysis:

(1) We identified FANTOM5 enhancers present only in a single cell type as follows: first, each enhancer, x, (out of n≈43k enhancers) is sorted by its signals across the cell types from highest to lowest. Second, a fold-change, FC(x), is computed between the first ranked cell type T_1_ and second ranked cell type T_2_. Lastly, given a threshold y, if FC(x) is above the y percentile of all n FCs, then enhancer x is considered as **uniquely active** (**ua**) in cell type T_1_ (termed **ua-enhancer**). We varied y between 40% and 95%. The higher the y threshold is - the more likely the enhancer that passed this criterion to be really active in a single cell type.

Notably, even for the y=95 percentile, the fold-change was a mere 2.3, very far from what could be considered a strictly unique cell-type enhancer. Moreover, with that threshold, only 12 enhancers were identified as uniquely active in GM12878.

(2) We took GM12878 POL2 ChIA-PET loops (m=95,269 loops) from (1) and resized the loop anchors to 5kb around their center position. We searched for loops whose anchors overlap with GM12878-ua enhancers. We counted how many ua-enhancers had an overlap with a loop anchor, and how many loops had anchors overlapping a ua-enhancer. We termed these loops as GM12878 specific E-loops (termed **sE-loops**).

For y=95, only 9 enhancers met this criterion, and they involved 45 sE-loops. So the vast majority of the experimentally obtained GM12878 loops (95,224/95,269 or 0.9995) did not involve a ua-enhancer, suggesting that this is a very rare case.

(3) Finally, we counted how many of the sE-loops involve anchors annotated as promoters (termed **sEP-loops**).

For y=95, only four loops met this criterion.

Even with the lowest tested threshold (y=40%, corresponding to a mere 1.0 fold-change), only 64 GM12878-ua enhancers were found. Out of them, 46 overlapped with 316 experimentally validated loops, and 70 loops involved an anchor annotated as promoter. So a truly unique enhancer is very uncommon.

We repeated the analysis for the cell type K562 using m=41,452 K562 ChIA-PET loops, and the numbers were even lower.

All the results are summarized in **Supplemental Table S3.**

Our analysis suggests that truly uniquely active enhancers are very rare. Therefore, we argue that it will be hard for any computational method to identify ct-links involving enhancers active in a single cell type.

We also tested a relaxed version of CT-FOCS that skips the leave-cell type-out cross validation step of FOCS and applies CT-FOCS to all 24,048 available promoters with their ten closest enhancers in FANTOM5. The latter method is termed 'CT-FOCS no filtering'. The reasoning was to allows a potential unique cell type not to be excluded in the cross validation. The results are shown in the table below. While CT-FOCS linked fewer promoters (~14K) than 'CT-FOCS no filtering' (~21K), both methods linked a similar number of enhancers (~27K). Also, both methods linked ~5K enhancers whose region appear in a single cell type (row 5 in the table – set A). Out of set A (row 6 in the table – set B), only ~200 enhancers (~4%) were ranked first by their signal in the same cell type compared to other cell types. The average and median FC (described in point 1 above) of the enhancers in set B was similar in both methods.

This analysis suggests that it is unlikely to observe enhancers that are strictly active in a single cell type. This may be because similar cell types from the same tissue have the same active enhancer. The more cell types are used in the analysis - the less likely we are to observe an enhancer active in a single tissue. Thus, the leave-cell-type-out-cross-validation step in FOCS has a minor effect on the identification of EP links that are strictly unique and active in a single cell type.

|  | CT-FOCS | CT-FOCS no filtering |
| --- | --- | --- |
| #candidate promoter models | 21,468 | 24,048 |
| #candidate enhancers | 36,244 | 37,193 |
| #linked promoters | 13,873 | 21,068 |
| #linked enhancers | 27,463 | 27,062 |
| A - #linked enhancers in a single cell type | 4,933 (18%) | 5,557 (20.5%) |
| B - #enhancers from A that were ranked first by signal in the same single cell type vs. other cell types | 196 (4%) | 175 (3.1%) |
| Avg/Median enhancer FC b/w first and second ranked cell types by enhancer signal (on the enhancers in B) | 1.5/1.3 | 1.8/1.2 |

**Supplemental Tables**

The supplemental tables are included in the online Supplemental Material in spreadsheet format. The 'Description' tab of each supplemental table file includes a table legend. Below are the titles of the tables:

**Supplemental Table S1:** Comparison between CT-FOCS, ABC model and TargetFinder on ENCODE DHS data across 5-10 cell types

**Supplemental Table S2:** The particularity of each algorithm's predictions as measured by ChIA-PET, HiChIP and PCHi-C assays

**Supplemental Table S3:** The number of ChIA-PET loops supported by enhancers with signal ranked above the y^th^ percentile, for different y values

**Supplemental Table S4:** FANTOM5 sample annotation. The annotation maps between 808 sample IDs, 472 cell types, and three cell type categories (cell line, primary cell, or tissue)

**Supplemental Figures**

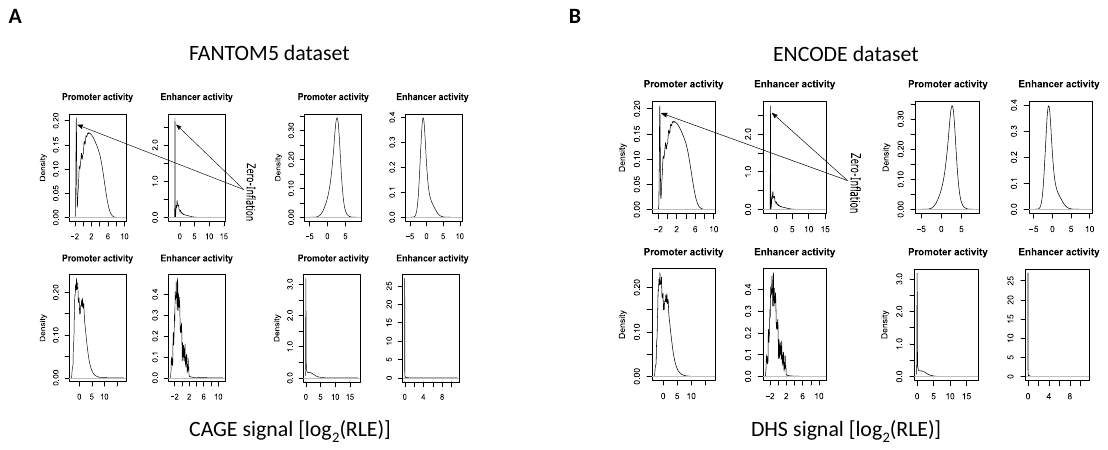

**Figure S1. Distribution of enhancer and promoter activity signals in FANTOM5 and ENCODE data.** **(A)** FANTOM5 (808 profiles) CAGE signal. The normalization of CAGE signals in CT-FOCS takes into account the library sizes. The activity of each enhancer or promoter was normalized using the RLE function (2) and log2 transformed. **(B)** ENCODE DHS signals (208 profiles). The sharp peaks in both promoter and enhancer indicate zero-inflated data.

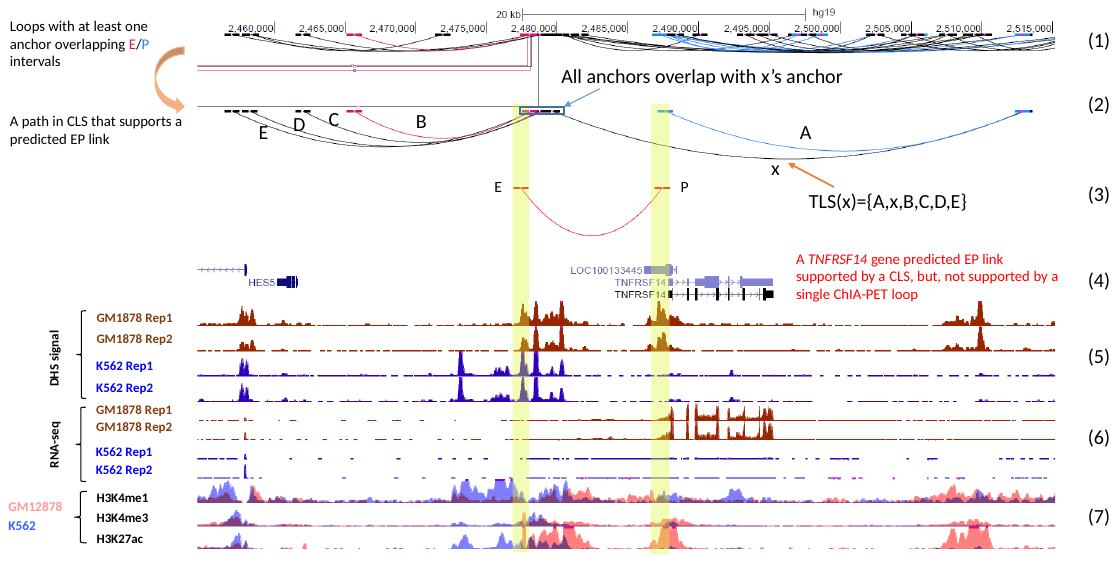

**Figure S2. ChIA-PET TLSs support ct-links.** TLS support for an EP link predicted in GM12878. From the top: (1) A track showing all ChIA-PET loops, highlighting those overlapping an enhancer (red) or a promoter (blue) in a segment of 60 kb of Chromosome 1. (2) All loops that have an anchor in common with loop x. All the anchors in the blue box have nonempty overlap. (3) The EP link predicted by CT-FOCS. That link is validated by TLS(x). (4) Gene annotation. (5) DHS signals of GM12878 and K562 (two replicates each). (6) RNA-seq levels of GM12878 and K562 (two replicates each). (7) Epigenetic marks of enhancer activity (H3K4me1+H3K27ac) and promoter activity (H3K4me3+H3K27ac). The TLS in the second track supports the single GM12878-specific EP link of *TNFRSF14* gene. This link is not supported by a single ChIA-PET loop. Tracks are shown using the UCSC genome browser.

| 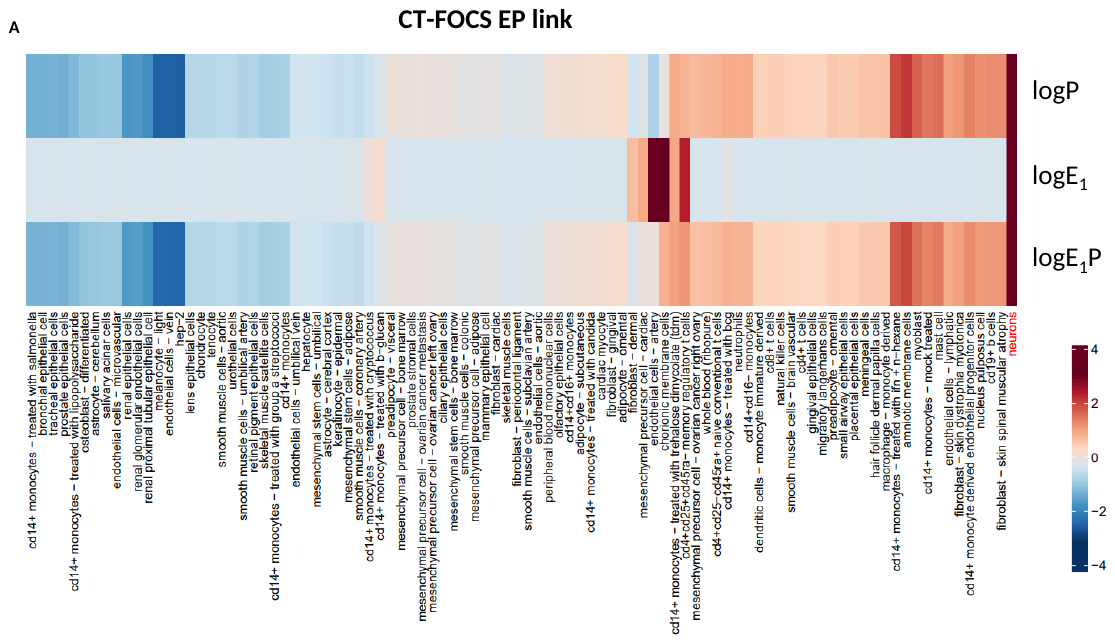 | |
| --- | --- |
| 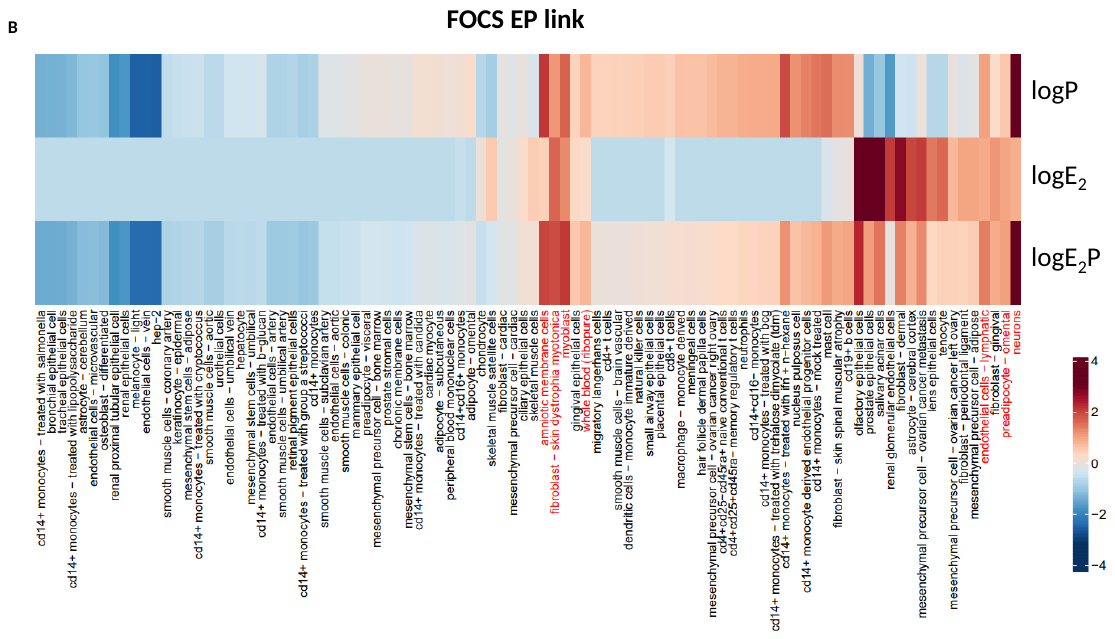 | |
| **Figure S3. The difference between CT-FOCS and FOCS in predicting EP links. (A-B)** Heatmaps of two EP links predicted by CT-FOCS and FOCS for the same promoter on FANTOM5 dataset. The links involve different enhancers, E_1_ and E_2_. **(A)** E_1_P link, predicted by CT-FOCS as specific for neurons primary cell. **(B)** E_2_P link predicted by FOCS. Cell type names marked in red have both enhancer and promoter signals above the 75% percentile across cell types. Only primary cells (n=94) with at least 3 replicates are presented. Values are in log_2_ median of the replicates per cell type. LogEP is the sum of logE and logP. For visibility, values in each row were transformed to the range -4 to 4. Samples were ordered by hierarchical clustering. | |
|  | 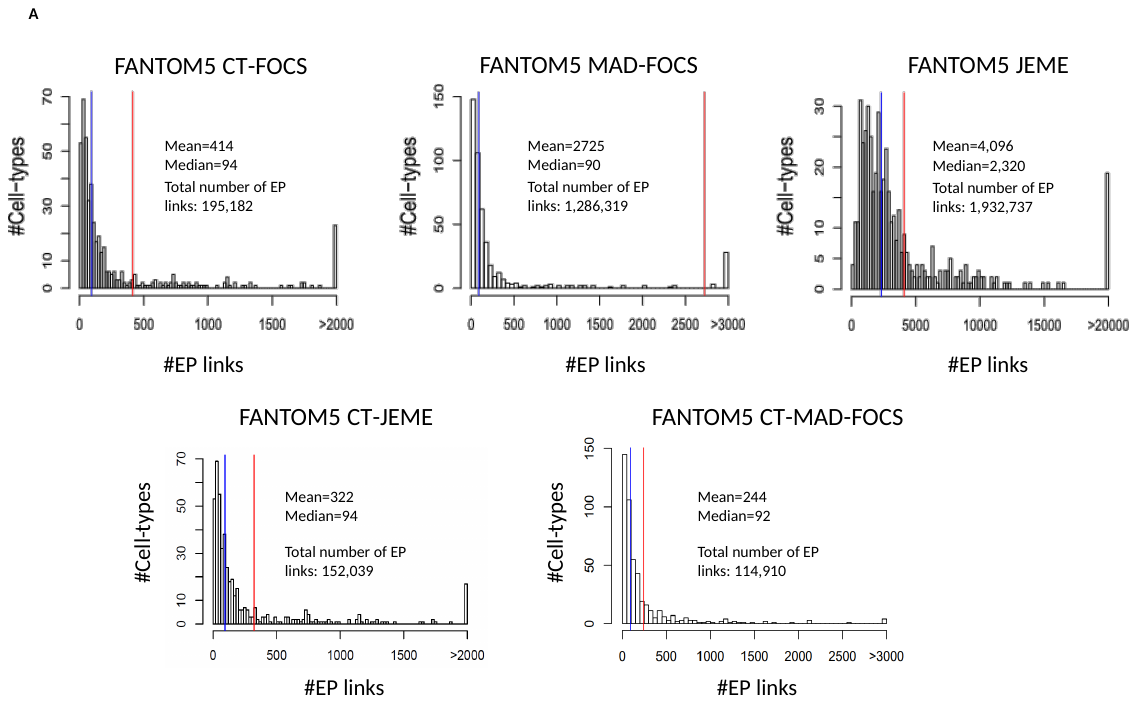  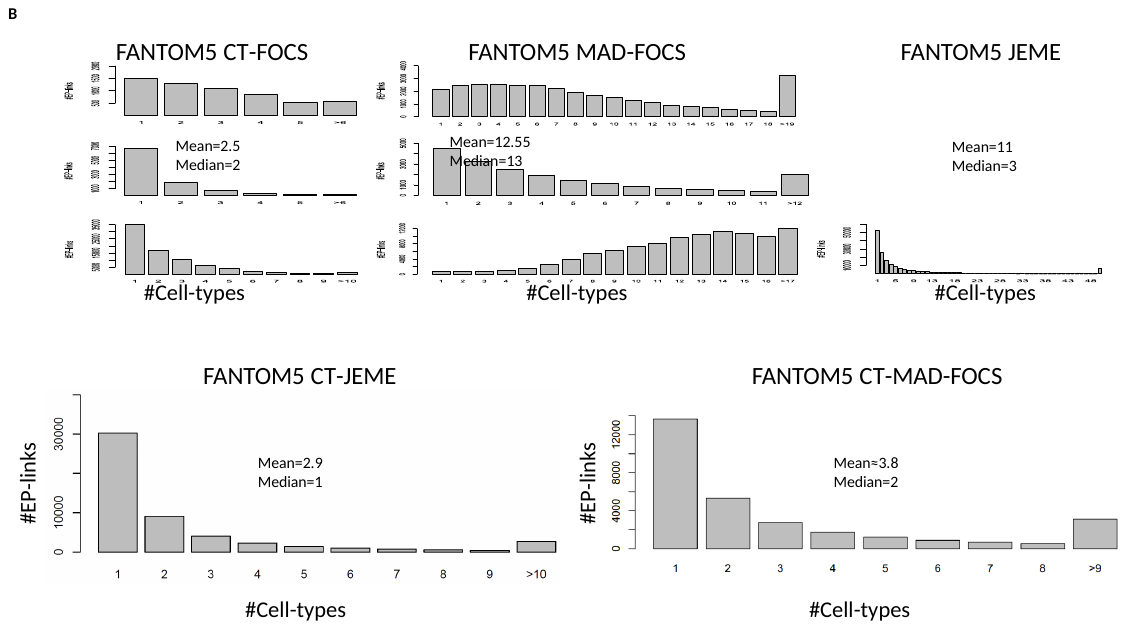 |
|  | 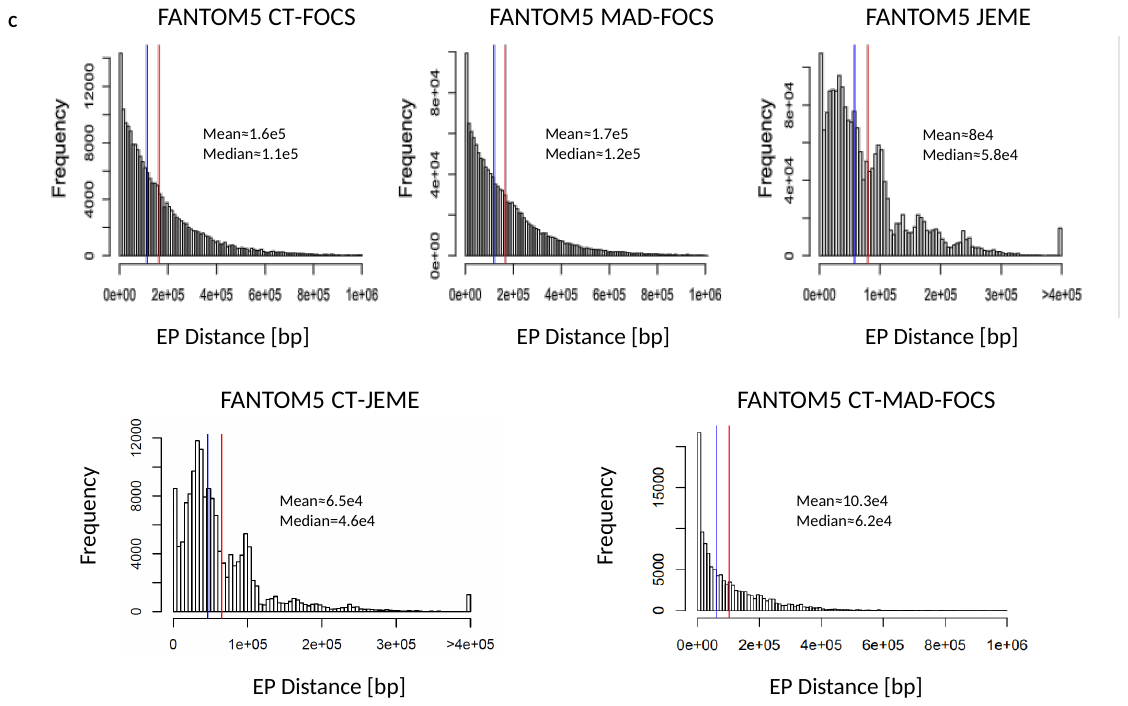 |
|  | **Figure S4. Properties of the ct-links predicted by CT-FOCS and by four other algorithms. (A)** Number of links predicted per cell type. **(B)** Sharing of ct-links among cell types. **(C)** Enhancer-Promoter distances in predicted ct-links. Distances were collapsed from all cell types, i.e., repeated EP links are counted multiple times. Predictions are on 472 cell types of the FANTOM5 data. |

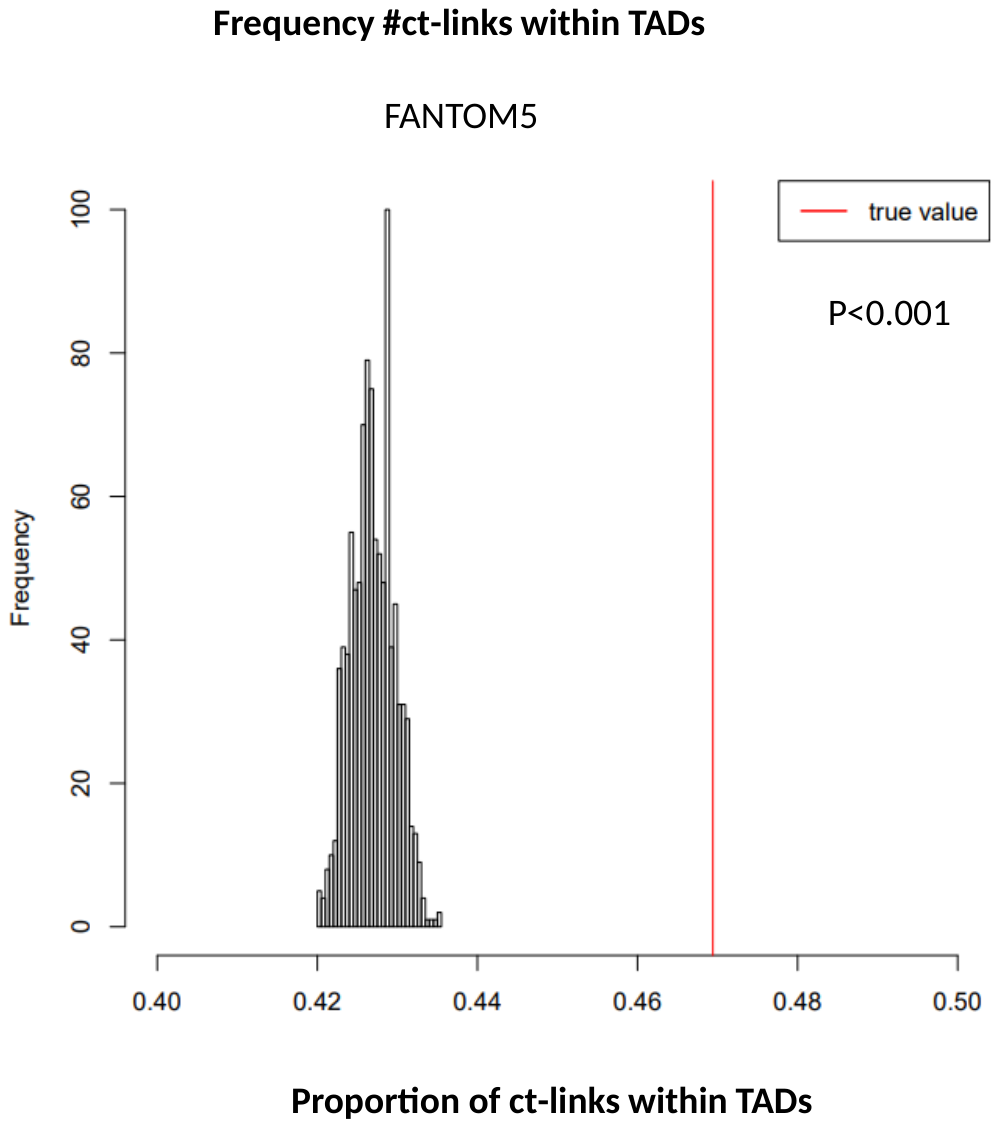

**Figure S5. Proportion of predicted ct-links that lie within TADs.** For each ct-link predicted by CT-FOCS, we randomly selected a promoter in the same chromosome and one of its 10 closest enhancers, and checked if they reside in the same TAD. The process were repeated 1,000 times, and the distribution of the fraction of links that fell within TADs is shown in black. The red line is the fraction obtained on the real data. The empirical p-value is the percentage of cases in which the fraction was higher than that observed on the real data (<0.001 here, as that percentage was zero). The 9,274 TADs reported in (3) were used.

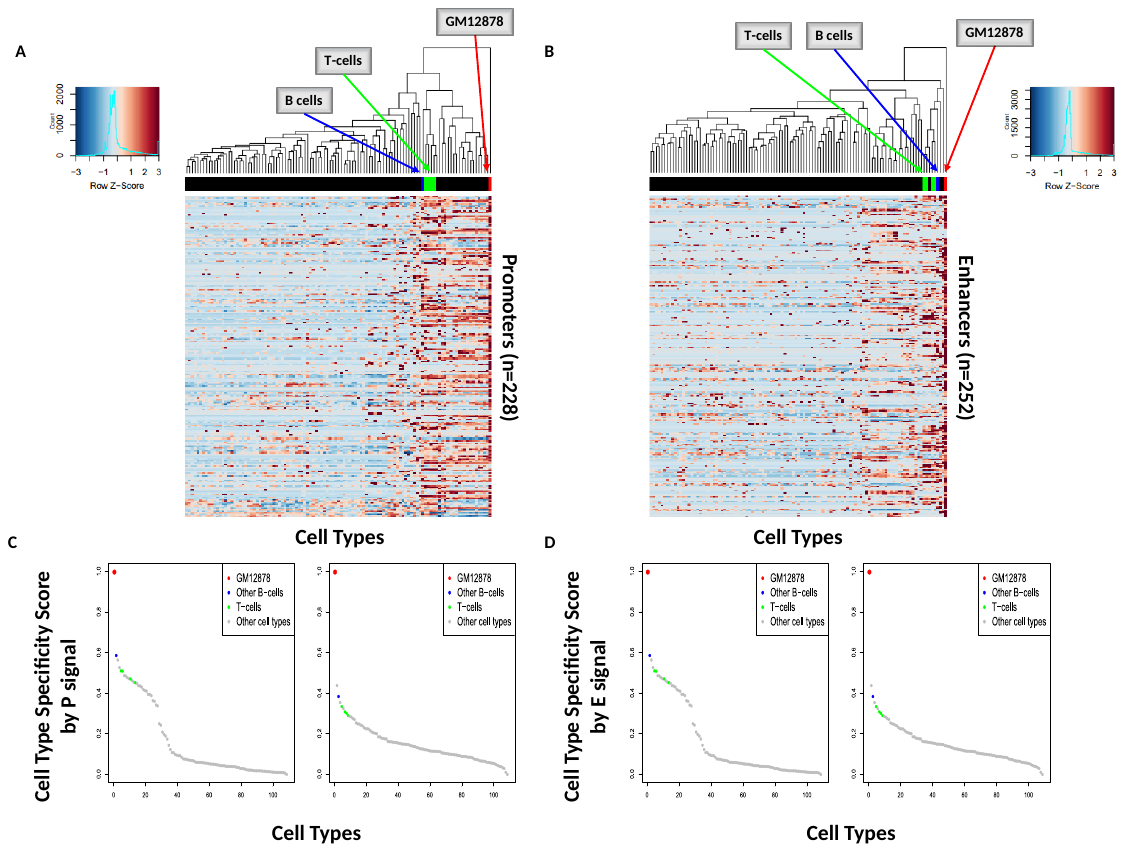

**Figure S6. Specificity of enhancers and promoters in ct-links predicted for GM12878. (A-B)** Heatmaps of linked promoter (A) and enhancer signals (B) for 340 ct-links predicted on GM12878. Columns – cell types, color – z-score of promoter and enhancer signal. B and T cell types related to GM12878 are highlighted in green and blue respectively. **(C-D)** Cell type specificity scores based on promoter (C) and enhancer signals (D). 109 cell types with at least 3 replicates each were included in the analysis (**Methods**).

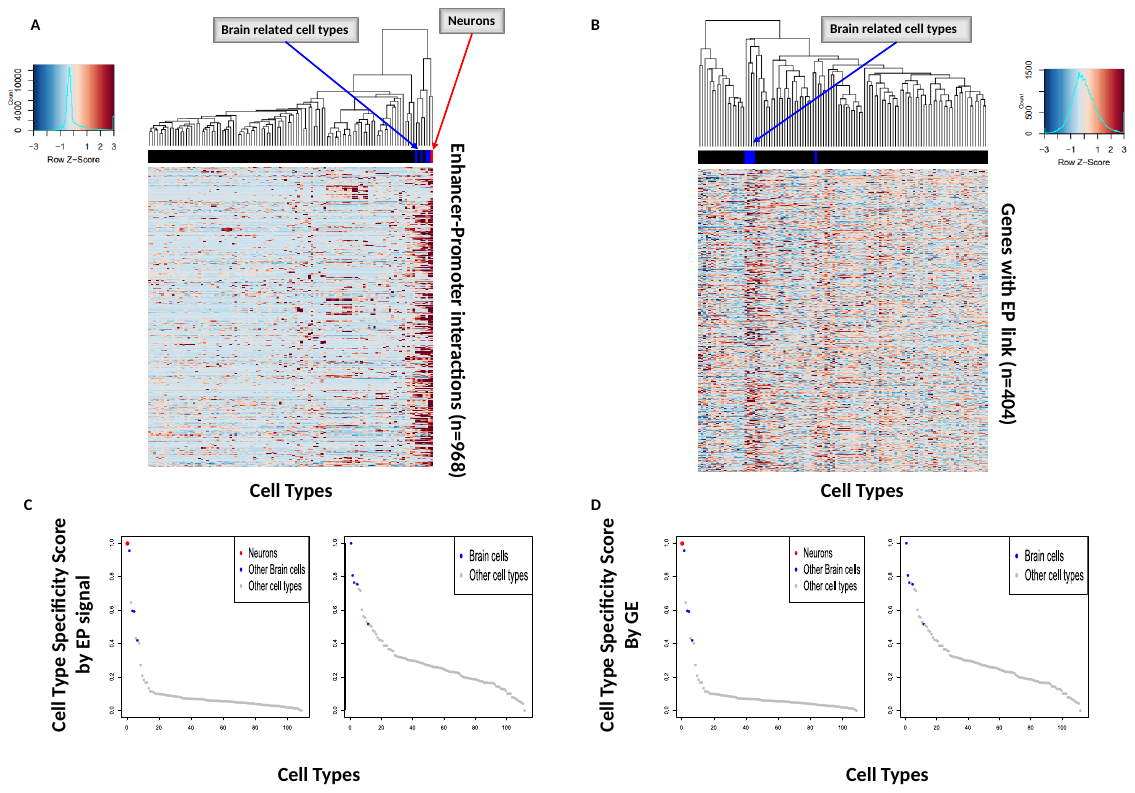

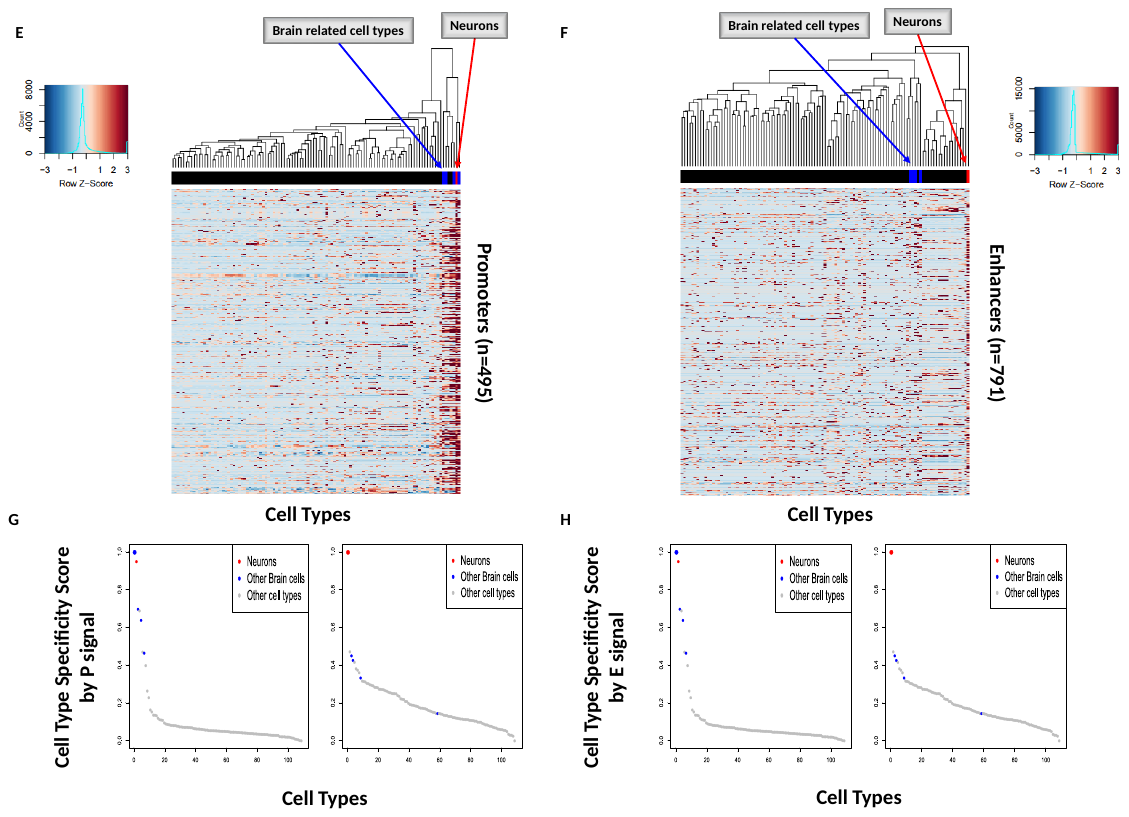

**Figure S7. Specificity of ct-links predicted for Neurons. (A)** Heatmap of EP signals for 968 ct-links predicted on Neurons based on FANTOM5 data. Rows – EP links, columns – cell types, color – z-score of EP signal. Brain cell types related to Neurons are highlighted in blue. **(B)** Heatmap of gene expression (GE) for 120 genes involved in the predicted ct-links. Rows – genes, columns – cell types, color – z-score of GE. **(C)** Cell type specificity scores based on the EP signals in A. **(D)** Cell type specificity scores based on expression for the gene set in B. **(E-F)** Heatmaps of linked promoter (E) and enhancer signals (F) for 968 ct-links predicted on Neurons. Columns – cell types, color – z-score of promoter and enhancer signal. **(G-H)** Cell type specificity scores based on promoter (G) and enhancer signals (H). In A, C and E-H, 109 cell types with at least 3 replicates each were included in the analysis; in B and D, 112 cell types with ENCODE gene expression are included (**Methods**).

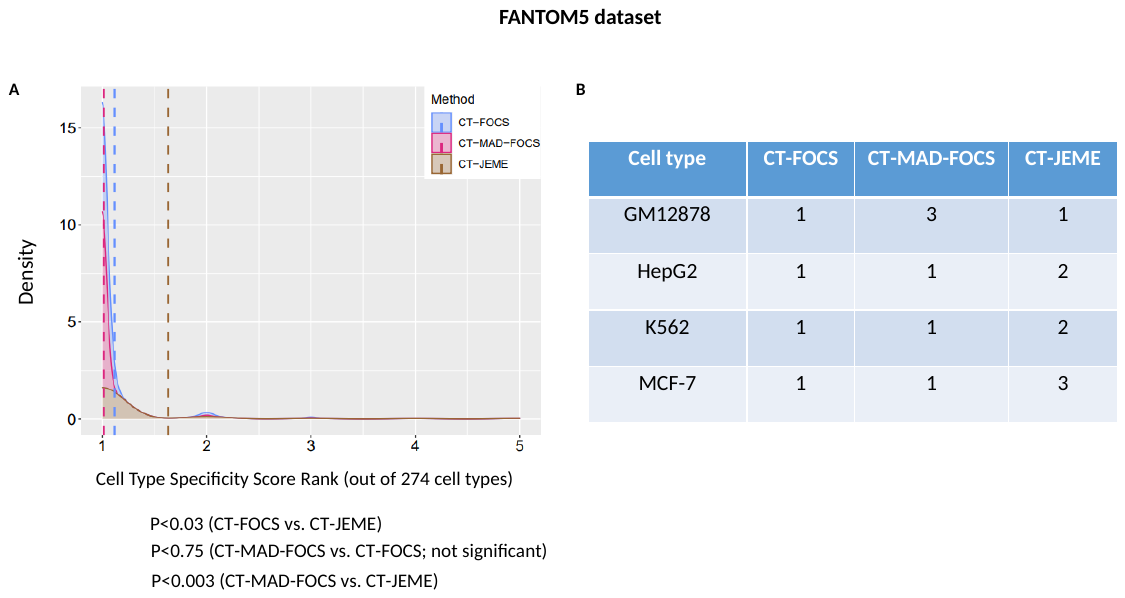

**Figure S8. Specificity scores of EP links predicted by CT-FOCS, CT-MAD-FOCS and CT-JEME. (A)** Density plots cell type specificity scores based on EP signals on 274 FANTOM cell types. Dashed lines denote the mean rank for each method. P-values were computed using one sided Kolmogorove-Smirnov test. **(B)** Ranking of the correct cell type in terms of specificity scores of the linked genes’ expression in four cell types for which expression data was available in ENCODE.

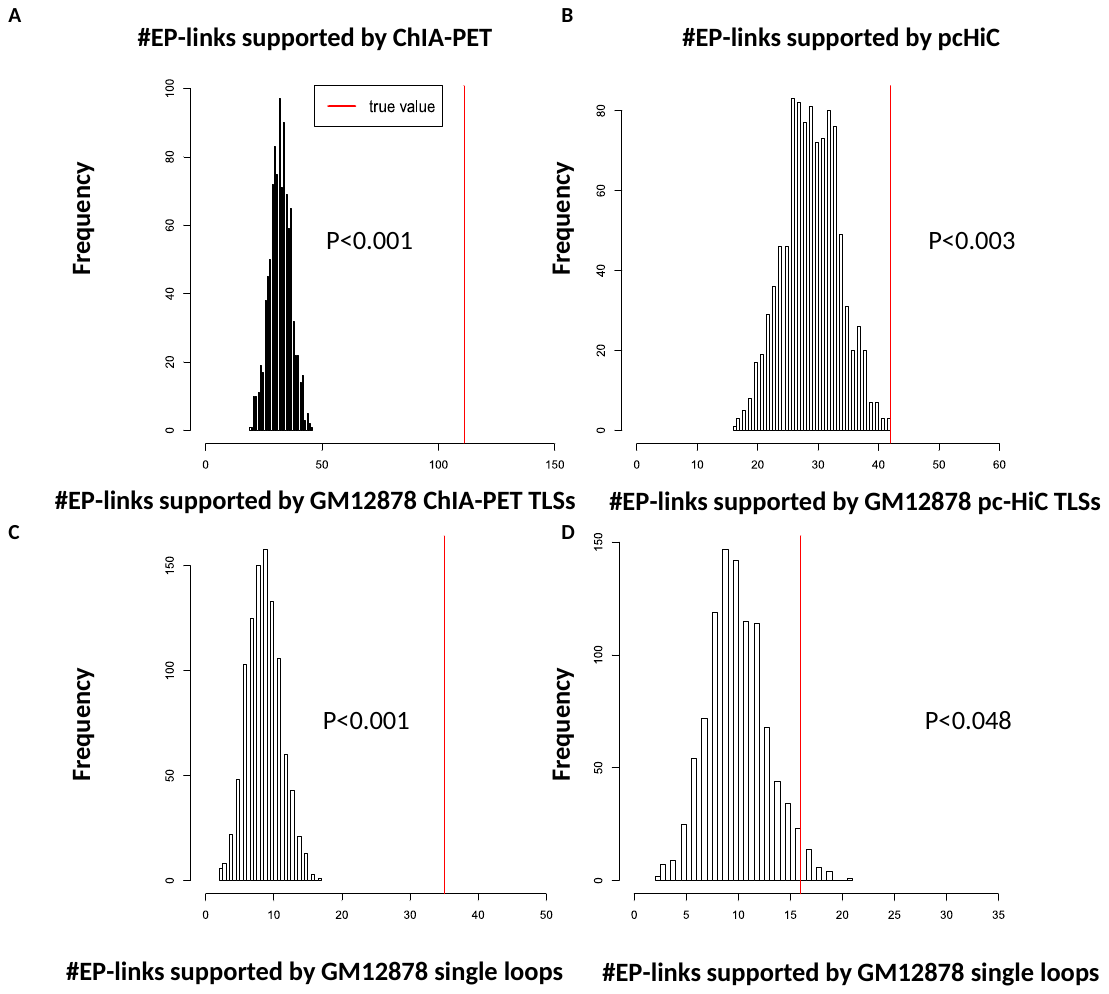

**Figure S9. Significance of the overlaps between predicted ct-links and experimental 3D contact data. (A-B)** Top – overlaps with GM12878 POL2 ChIA-PET (A) and pcHiC (B) TLSs. **(C-D)** Bottom - overlaps with GM12878 ChIA-PET (C) and pc-HiC (D) single loops. Random sets of EP links of the same number and linear distance as the true ct-links were generated. In each set, the number of random EP links that overlapped with the TLSs or single loops in 3D data was counted. The black distribution shows the counts for 1000 random sets, and the red line shows the number obtained for the links inferred by CT-FOCS. Except (D), in all cases the number of random sets with higher counts than the true set was zero.

**Supplemental Methods**

**ENCODE DHS data preprocessing**

ENCODE DNase-seq samples (106 cell types) were downloaded from GEO dataset GSE29692 (4–6). ENCODE DHS peaks of enhancers and promoters (7) were processed as in FOCS (8) with the following changes: (1) we analyzed only promoters of annotated protein-coding genes according to GencodeV10 TSS annotations (<ftp://genome.crg.es/pub/Encode/data_analysis/TSS/Gencodev10_TSS_May2012.gff.gz>). (2) We applied a relative-log-expression (RLE) normalization (2), as implemented in edgeR (9, 10). (3) We retained promoters and enhancers that showed robust activity in at least one cell type: signal ≥ 5 RPKM in all samples of at least one cell type. Overall, we analyzed 208 samples from 106 cell types. Our preprocessing resulted with 36,056 promoters (mapped to 13,105, 13,464, and 13,197 protein-coding genes according to HGNC_symbols, Ensembl, and Entrez, respectively) and 658,231 putative enhancers.

Enhancers closer than 10kb to the nearest promoter were discarded since we wanted to reduce false positive links due to the high signal correlation at short distances, and to predict distal interactions as suggested by Whalen et al. 2016. The candidate enhancers for each promoter were defined as the 10 closest enhancers located within a window of 1Mb (±500kb upstream/downstream) from the promoter’s center position.

We first applied the FOCS pipeline, including leave-cell-type-out cross validation (LCTO CV) on the promoters and their candidate enhancers, and accepted promoter models with q-value$\leq0.1$ in the activity level test (see Hait et al. 2018 for details). Unlike FOCS, we did not apply here regularization on the predicted EP links. Overall, the procedure resulted with 17,832 promoter models (mapped to 9,090, 9,320, and 9,160 HGNC_symbols, Ensembl, and Entrez protein-coding genes, respectively).

**FANTOM5 CAGE data preprocessing**

We downloaded the FANTOM5 CAGE data from JEME (12) repository (<https://www.dropbox.com/sh/wjyqyog3p5d33kh/AACx5qgwRPIij44ImnzvpFxUa/Input%20files/FANTOM5/1_first_step_modeling?dl=0&subfolder_nav_tracking=1>). Overall, the data contained 24,048 promoters (mapped to 18,986, 20,597 and 18,912 protein-coding genes according to HGNC_symbols, Ensembl and Entrez, respectively) and 42,656 enhancers, covering 808 samples. Enhancer and promoter expression matrices were RLE normalized. We used the RLE normalized data from (12). We manually annotated the 808 samples with 472 cell types (**Supplemental Table S4**) using **Table S1** from FANTOM5 (13).

For each promoter, the candidate enhancers were defined as the 10 closest enhancers located within ±1Mb from the promoter’s TSS as performed in JEME (12). Unlike ENCODE, we did not enforce a lower bound on the distance here. We applied the same pipeline on the promoters and their candidate enhancers as described above for the ENCODE data. This resulted with 21,468 promoter models for further analysis.
